# Assessment of neurobehavioural traits under axenic conditions: an approach for multiple longitudinal analyses in the same mouse

**DOI:** 10.1101/2024.11.21.623170

**Authors:** Andrina Rutsch, Monica Iachizzi, Jorum Kirundi, Johan B. Kantsjö, Arielle L. Planchette, Terry Müller, Werner Schmitz, Aleksandra Radenovic, Raphaël Doenlen, Mercedes Gomez de Agüero, Francesca Ronchi

## Abstract

The gut-microbiota-brain axis influences neuroinflammation, neural development and behaviour such as sociability, memory and anxiety. To study these traits *in vivo*, especially during development or disease, it is crucial to analyse them over time and with multiple analyses in the same animal. With a growing understanding of the role of specific bacteria in neurodegenerative disease and behaviour, the demand for gnotobiotic mouse models has increased. However, maintaining stable hygienic conditions during behavioural testing is challenging, as exposure to conventional environments can alter the hygienic status of mice and affect behaviour. We established protocols to perform behavioural tests assessing memory, anxiety, exploration, learning and recognition under axenic conditions using flexible film isolators. Our study compared the behaviour of germ-free mice with mice carrying a defined minimal or moderately diverse microbiota. The results showed no effect of the microbiota on short- and long-term memory or novel object recognition. However, we showed for the first time that mice colonised with defined commensal bacteria exhibited more anxiety-like behaviour than germ-free mice. In addition, we showed that microbiota complexity is important, as only mice colonised with moderately diverse microbiota exhibited anxiety-like behaviour, allowing us to disentangle the contribution of specific microbial species or community interactions to this phenotype. This phenotype correlated with differences in hippocampal and serum metabolic profiles between colonised and germ-free mice. We propose a novel approach to study rodent behaviour at different physiological and pathological stages in their life without compromising hygiene, thus promoting the refinement and reduction of mice used in experiments.

## Introduction

The gut-microbiota-brain (GMB) axis is the multidirectional communication between the gut, its microbiota and the central nervous system (CNS). Increasing evidence suggests that cognitive changes are influenced by the composition of the gut microbiota^1–3^. Gnotobiology and defined microbial consortia^4–9^ have become valuable tools for studying the impact of the microbiota on health and disease^9^. Consequently, there is a growing need for publicly available, stable and reproducible gnotobiotic mouse models to study their behaviour. A challenge in behavioural testing under gnotobiotic conditions is to avoid exposure of the animals to the environment, which can affect their hygiene status and subsequentially the outcome of the tests^10^. Recent studies^10–12^ have developed systems to assess the behaviour of laboratory rodents in a sterile isolator, focusing on tests such as light/dark preference, step-down, tail suspension and marble burying. These tests are widely used to respectively assess anxiety-like behaviour, aversive memory and learning, depressive-like behaviour and anxiety or obsessive-compulsive behaviour. These studies have shown that colonisation with either diverse (conventionally housed) bacteria, highly restricted microbiota (Altered Schaedler Flora, ASF^13,14^), a single bacterial strain (*Escherichia coli* JM83) and even a transient colonisation with an auxotrophic strain of *E. coli*, can normalise exploratory/anxiety-like behaviour and brain metabolome of germ-free (GF) mice^12^. In several reports, ASF microbiota promoted anxiety-like behaviour in mice with higher corticosterone levels than in conventional mice^15^. Despite these findings, the gnotobiotic models tested in these studies, such as ASF consortia or monocolonisation models, may not fully recapitulate the physiology of conventionally colonised animals^14,16,17^.

To fill this gap, we studied behaviour in a stable, defined, moderately diverse microbiota model called Oligo-MM^12^ ^4–6,18^, which better mimics the physiology of highly diverse colonised mice than the highly reduced ASF model^19^. We compared Oligo-MM^12^ mice with age- and sex-matched GF mice and those colonised with a minimal (four bacteria, called Cuatro^20^) microbiota. For the first time to our knowledge, we have introduced tests to assess different aspects of learning and memory^21^ in mice, such as short- and long-term memory, anxiety and exploration, learning and recognition while maintaining their hygiene status, in sterile flexible film isolators, allowing for longitudinal studies. Our results showed that Oligo-MM^12^ ^4–6,18^ mice exhibited more anxiety-like behaviour than GF or Cuatro mice, while no significant differences were found in memory or recognition abilities. In addition, housing conditions influenced the results, with notable differences between isolators and conventional facilities. We also found differences in hippocampal and serum metabolic profiles between our gnotobiotic mouse models, suggesting a correlation between microbiota-dependent expression of neuroactive molecules and behaviour. For the first time, we have characterised multiple behavioural traits of Oligo-MM^12^ gnotobiotic mice and propose a methodology that can be easily adopted in animal facilities and experimental laboratories to assess different brain functions in the same animal over time, while maintaining a stable hygienic condition during physiological and pathological development and reducing the number of experimental mice used.

## Results

### Sequential behavourial tests under axenic conditions

To perform sequential behavourial tests under controlled hygienic conditions, behaviour testing equipment was installed in a sterile flexible film isolator to maintain and control the hygienic status of the mice throughout the tests (**Fig. S1A**). The mice underwent sequential tests as follows: first the short-term memory (STM) test (week 1), the long-term memory (LTM) test (week 2), then the open field test (week 3) followed by the novel object recognition test (week 3) (**Fig. 1A**), to analyse spatial and non-spatial learning, memory and anxiety-like behaviour respectively under axenic conditions.

**Fig. 1:**
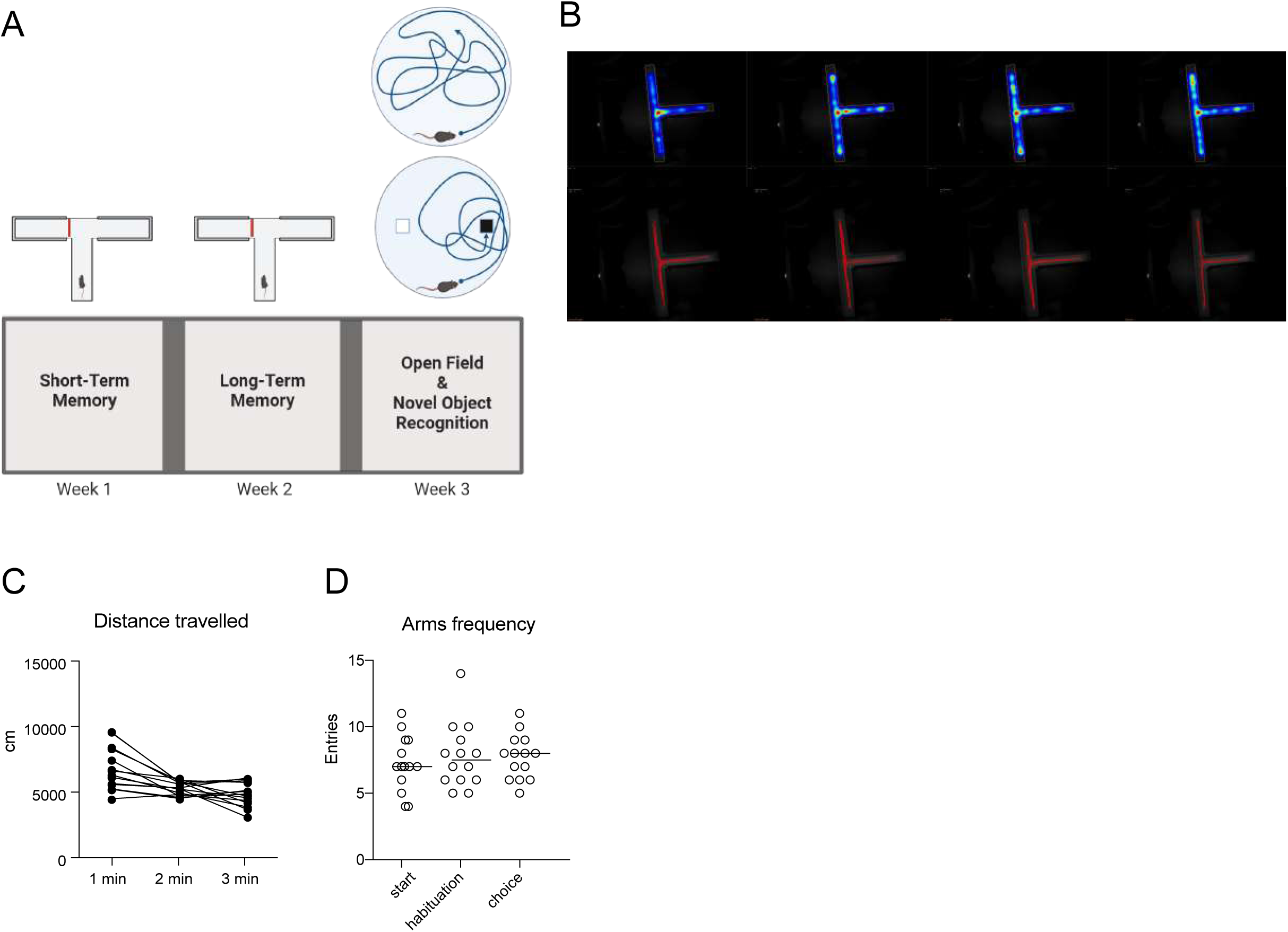
Experimental strategy of the different tests under axenic conditions. Various behavioural tests under axenic conditions inside sterile flexible film isolators. (**A**) Sequential behavourial under axenic conditions: short-term memory (STM) test first (week 1), the long-term memory (LTM) test (week 2), the open field test (week 3) followed by the novel object recognition test (week 3) (**B**) Heat maps (top) and tracks (bottom) of the movement of the mice (Adult female C57BL/6 germ-free (GF) mice shown) during the three minutes of exploration in the T-maze arena. (**C**) Total distance travelled (cm) and (**D**) number of entries into the different zones of the arena during the entire duration of exploration of the T-maze arena. The mean is indicated by the bar height with standard deviation. Each point represents an individual mouse. Sample size: n = 14.

STM and LTM tests were performed in the T-maze arena to assess short- and long-term spatial working memory in mice (**Fig. S1B-F**). The T-maze consists of a stem (starting arm) and two lateral arms, the left (choice) and right (habituation) arms (**Fig. S1B, C**).

To exclude any external confounding factors, such as light, shadows and reflections on the arena walls as well as operator manipulation, we conducted a pilot experiment with adult GF mice. These mice freely explored the T-maze arena for three minutes and their preference for certain parts of the T-maze was recorded (**Fig. 1B-D**). The mice explored the arena homogeneously, being more active in the first minute of exploration (**Fig. 1B, C**) and showed no arm preference throughout the test (**Fig. 1D**). This confirmed that our experimental conditions within the isolator were appropriate for the established settings.

### Microbiota does not shape short and long-term memory under axenic conditions

To test the effect of microbiota diversity on spatial learning and memory function, we performed short-term and long-term memory tests in the T-maze arena in gnotobiotic mice with a moderately diverse microbiota (Oligo-MM^12^ model) and compared them with GF and Cuatro mice. During four consecutive habituation trials, the mice were allowed to freely explore the start and habituation arms for two minutes each trial, followed by one minute or 24 hours rest for STM or LTM, respectively, to familiarise themselves with the environment, while the choice arm was closed (**Fig. S1D, E**). In trial 5 (choice), the mice were free to explore the entire T-maze for two minutes, including the unfamiliar arm for the first time. The number of entries and duration in each arm was analysed. This test takes advantage of the rodents’ natural tendency to explore unfamiliar areas of the environment rather than familiar ones (novelty preference). To discriminate between familiar and unfamiliar arms in trial 5, the animal must spatially remember which arm it visited, which arm was open in the previous trials and which arm is unfamiliar. In both STM and LTM (**Fig. 2, Fig. S1F**), GF and Cuatro animals behaved similarly in terms of distance travelled and number of entries in both the start and habituation arms throughout the duration of the tests, irrespective of their hygiene status (**Fig. 2A-F**). Oligo-MM^12^ mice travelled less distance and entered the start arm less often during the choice phase in the STM test. During the different habituation phases in the LTM test, they entered less often in the start and habituation arms (**Fig. 2A, E, F**). Indeed, the proportion of Oligo-MM^12^ mice entering the choice arm at least once was lower than in GF or Cuatro mice (**Fig. 2I**). This could indicate that mice colonised with more diverse bacteria are more reluctant to explore a change in a familiar environment, which could also be reflected in their shorter distance travelled during the choice phase in STM (**Fig. 2A**). However, there was no significant difference in the number of entries or the time spent in the choice arm for this group of mice compared to the others (**Fig. 2G-H**), and the tendency was also less pronounced in the LTM test (**Fig. 2J-L**). These data showed that the microbiota does not shape short- and long-term memory.

**Fig. 2:**
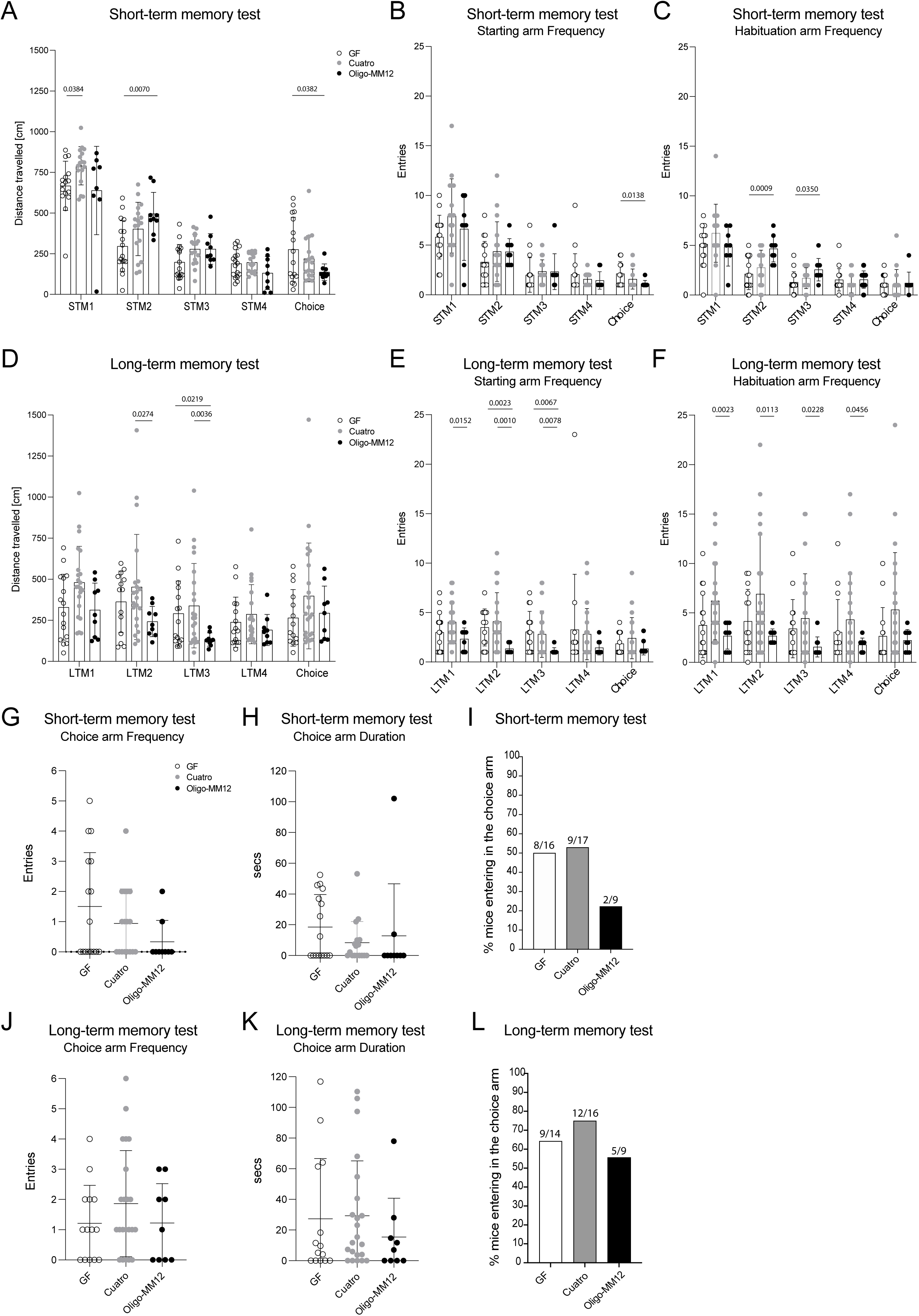
Microbiota does not shape short and long-term memory. Adult female C57BL/6 mice, born and raised under germ-free (GF) or gnotobiotic conditions (Cuatro or Oligo-MM^12^), were subjected to Short-Term (STM) and Long-Term Memory (LTM) tests in the T-Maze at the Clean Mouse Facility of the University of Bern. (**A-C, G-I**) Short-Term and (**D-F, J-L**) Long-Term Memory test. (**A, D**) Total distance (cm) travelled in STM (**A**) and LTM (**D**) during the entire duration of each test in the different indicated habituation sessions. (**B, C, G; E, F, J**) Number of entries in each different arm (**B, E**, starting arm; **C, F**, habituation arm; **G, J**, choice arm) of the T-maze during the entire duration of the test. (**H, K**) Time (secs) spent in the choice arm during the choice session. (**I, L**) Number and frequencies of mice that entered the choice arm at least once during the STM (**I**) and LTM (**L**). Mean indicated by bar height with standard deviation. P values are indicated in the graphs and determined by one-way ANOVA followed by Tukey’s multiple comparisons test. Each dot represents a single animal. Sample size: n ≥ 9.

### Microbial diversity affects anxiety-like behaviour

In week 2, the same animals were tested in the open field test (**Fig.S1A; S2; S3A, B**) to measure locomotion, anxiety-like and exploratory behaviour. Mice showing more anxiety-like behaviour spend more time near the walls of the arena, in the periphery, as opposed to the centre of the arena. Mice were recorded freely exploring the novel environment for 20 min (**Fig. S3A**). Oligo-MM^12^ mice preferred to spend more time in the periphery of the arena, rather than in the centre, compared to GF or Cuatro mice (**Fig. 3A**). However, all mice travelled a similar distance over the duration of the experiment under all different hygiene conditions, showing that the locomotor activity was not affected by the hygiene condition (**Fig. 3A, B**). This suggests that Oligo-MM^12^ mice exhibited more anxiety-like behaviour compared to GF or Cuatro mice, confirming previous findings comparing GF mice and mice under different hygiene conditions^12,15,22,23^. This characteristic may also be reflected in the tendency of Oligo-MM^12^ mice to enter the different arms of the T-maze arena less frequently during the LTM test (**Fig. 2E, F**). To understand whether the different behaviour of the Oligo-MM^12^ mice observed under axenic experimental conditions was also reproducible under experimental conditions typical of conventional animal behaviour testing facilities, additional GF and Oligo-MM^12^ colonised mice were shipped from the University of Bern to the Center of Phenogenomics in Lausanne (EPFL) (**Fig. 3C, D**) and tested together with age- and sex-matched SPF mice. Even under classical experimental conditions, where GF and Oligo-MM^12^ colonised mice are exposed to the environment and their hygiene status is compromised, Oligo-MM^12^ and GF mice tested at the EPFL showed anxiety-like behaviour similar to the mice of the same hygiene status tested at the University of Bern, with Oligo-MM^12^ colonised mice spending more time in the periphery of the arena than the GF animals (**Fig. 3C, D**). Interestingly, the Oligo-MM^12^ mice showed anxiety-like behaviour similar to that of the SPF mice. Notably, the arenas used in Bern and in Lausanne were different in size, suggesting that neither arena size nor environmental conditions affected the open field behaviour in Oligo-MM^12^ mice as previously suggested^24^. In conclusion, our data indicate that the open field test, performed in flexible film isolators under axenic conditions, is a valid tool for testing anxiety-like behaviour in mice.

**Fig. 3:**
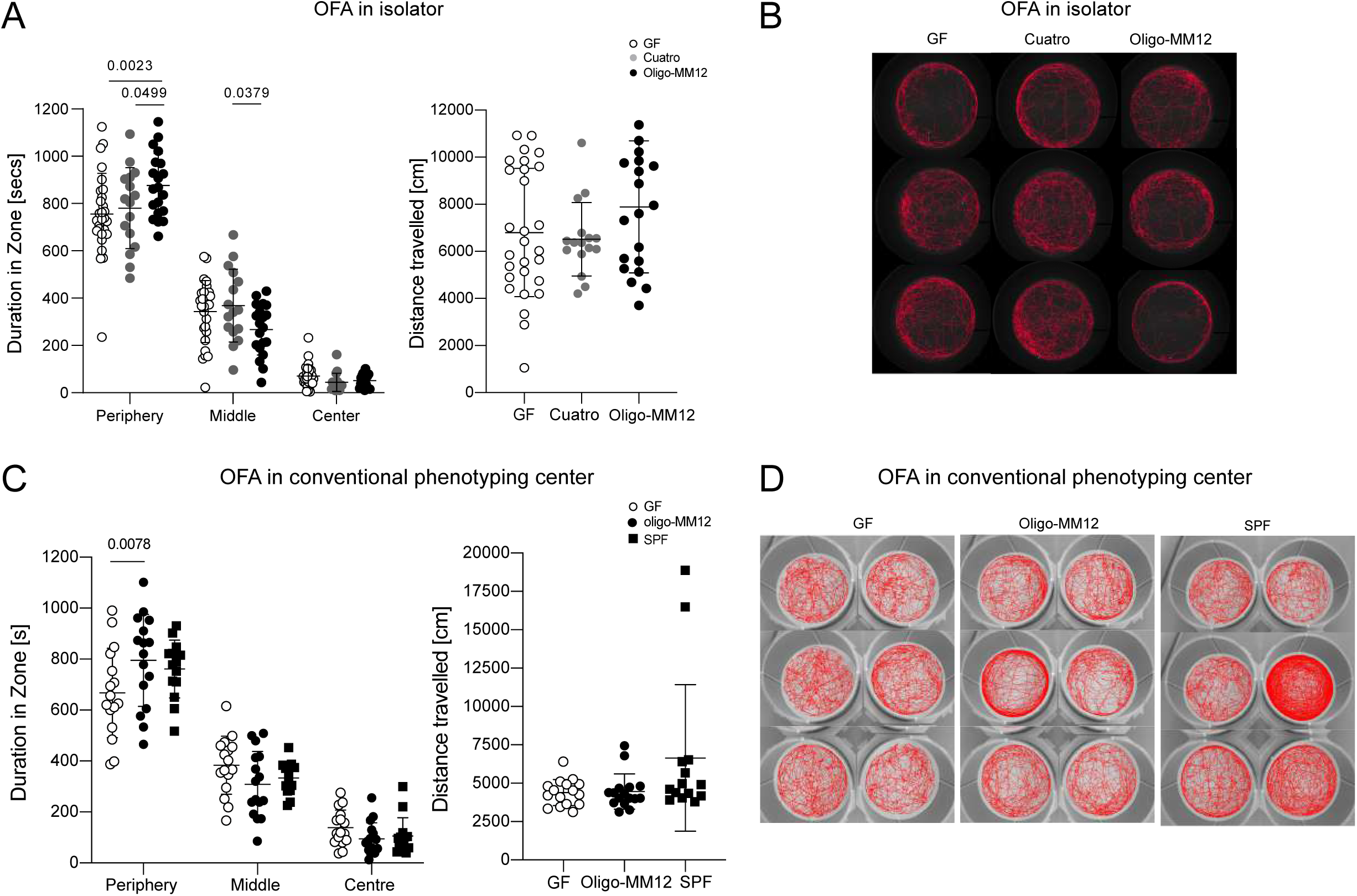
Oligo-MM^12^ colonised mice show more anxiety-like behaviour than germ-free or Cuatro-colonised mice. (**A-D**) Adult female C57BL/6 mice, born and reared under GF or gnotobiotic conditions (Cuatro and Oligo-MM^12^), were subjected to an Open Field Test in two different animal facilities under different hygiene conditions. (**A, B**) Tests performed at the Clean Mouse Facility of the University of Bern under axenic conditions, in sterile flexible film isolators. (**A**) Duration (s) spent in the different indicated zones of the arena for the whole test (left graph) and total distance travelled (cm) during the whole test (right graph). (**B**) Tracks of the animals’ movements during the whole test, each track picture is a different mouse. (**C, D**) Tests performed at the Center of Phenogenomics of the EPFL in Lausanne under conventional conditions in a small arena. (**C**) Duration (s) spent in the different indicated zones of the arena during the whole test (left graph) and total distance travelled (cm) during the whole test (right graph). (**D**) Tracks of the animals’ movements during the whole test, each track picture is a different mouse. Mean is indicated by bar height with standard deviation. P-values are indicated in the graphs and determined by one-way ANOVA followed by Tukey’s multiple comparison test. Each point represents a single animal. Sample size: n ≥ 9.

### The ability to recognise a new object in a known environment is highly dependent on the housing and testing conditions rather than the microbiota

Following the open field test, the same mice were tested for cognition, specifically for recognition memory, using the Novel Object Recognition test, both in axenic conditions (at the University of Bern) and in conventional environments (at the EPFL) (**Fig. 4, Fig.S3C, D**). During the first (learning) phase of the Novel Object Recognition test, mice are exposed to two identical objects in the open field arena. In the second (recognition) phase, one of the objects is replaced by a novel object of the same size but a different colour. The test assesses the mouse’s ability to recognise and explore the new object, again exploiting the rodent’s natural preference for novelty. The amount of time spent exploring the novel object compared to the time spent exploring the familiar object over the total object exploration time provides an index of recognition memory (recognition index = RI). All mice travelled a greater distance during the learning phase than during the recognition phase (**Fig.4A, C**). In the isolator, GF and Cuatro mice showed a significant ability to recognise a novel object (NO) over a familiar object (FO1). Oligo-MM^12^ mice showed a tendency to recognise a novel object (NO) over a familiar one (FO1), but this was not statistically significant (**Fig.4B**). Interestingly, in the environment of the conventional phenotyping center at EPFL, Oligo-MM^12^ mice showed more interest in the novel object (NO) than the familiar one (FO1), in contrast to GF mice (**Fig.4D**), which showed no preference for the NO in this environment, similar to what has been previously reported for SPF mice compared to antibiotic-treated animals^25^.

**Fig. 4:**
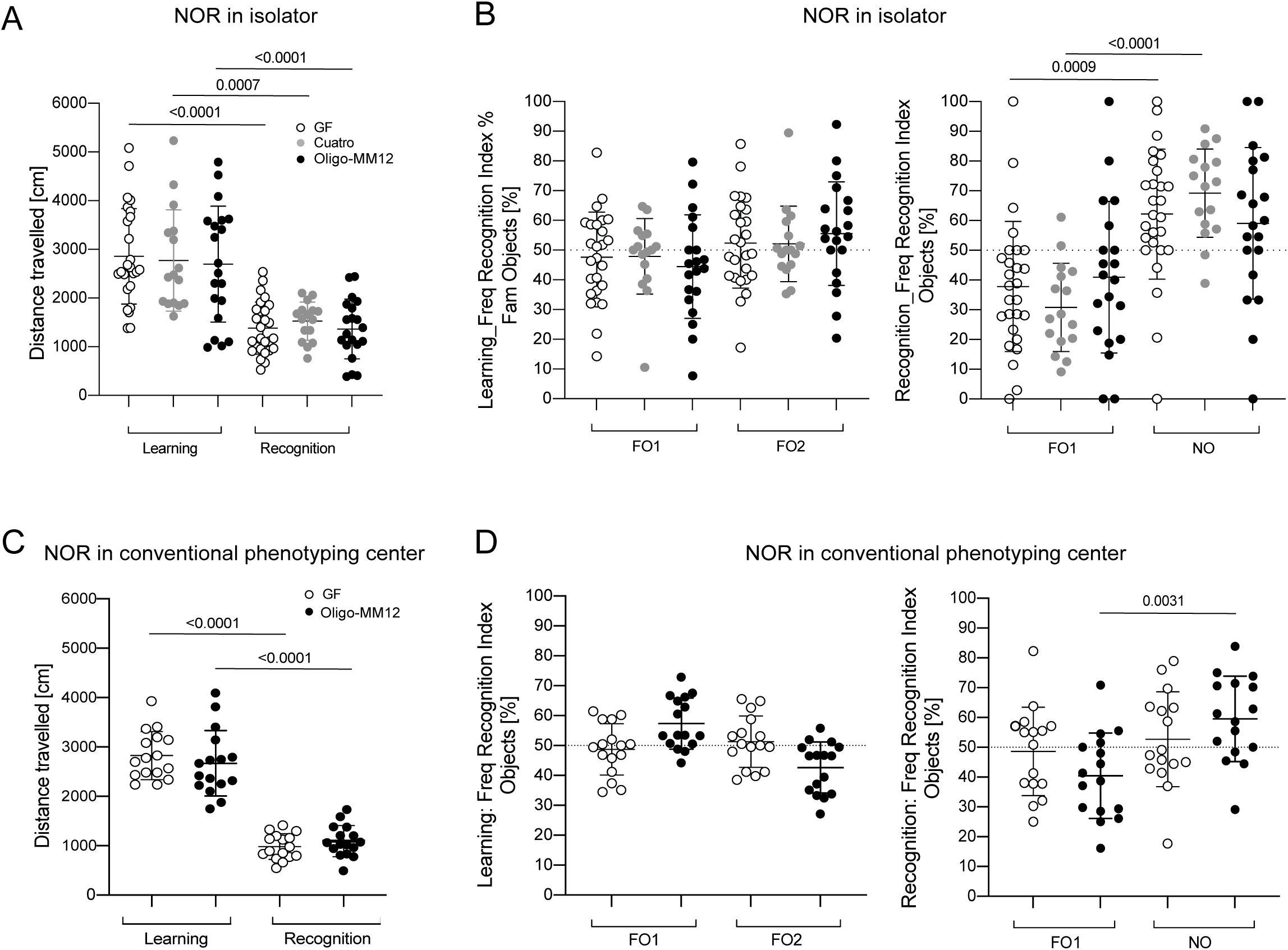
The ability to recognise a new object in a known environment is highly dependent on the housing and testing conditions rather than the microbiota. Adult female C57BL/6 mice, born and raised under GF or gnotobiotic conditions (Cuatro and Oligo-MM^12^), underwent Novel Object Recognition test in two different animal facilities under different hygiene conditions. (**A, B**) Tests performed at the Clean Mouse Facility of the University of Bern under axenic conditions, in sterile flexible film isolators. (**C, D**) Tests performed at the Center of Phenogenomics of the EPFL in Lausanne under conventional conditions in a small arena. (**A, C**) Total distance travelled (cm) in the learning phase and in the recognition phase, during the entire duration of the test. (**B, D**) Frequency Recognition Index (%) for both objects during the learning phase (left) and during the recognition phase (right). The index Is calculated as the ratio of the number of times the mouse explored the relevant object (exploration = entry into object zone while nose point facing the object) and the total number of times the mouse explored both objects. Mean with standard deviation are indicated. P values are indicated in the graphs and determined by one-way ANOVA followed by Tukey’s multiple comparisons test. Each dot represents a single animal. Sample size: n ≥ 9. Mean is indicated by bar height with standard deviation. P-values are indicated in the graphs and determined by one-way ANOVA followed by Tukey’s multiple comparison test. Each point represents a single animal. Sample size: n ≥ 9. FO1 = Familiar Object 1, FO2 = Familiar Object 2, NO = Novel Object.

These results suggest that the environmental/housing conditions (e.g. flexible film sterile isolator or conventional settings) significantly affect the novel object recognition ability of mice. Therefore, careful consideration should be given when designing experiments and selecting methods to assess biological parameters^26^. However, in both conditions, Oligo-MM^12^-colonised mice show different behaviour than GF or Cuatro mice.

### Oligo-MM^12^ microbiota recapitulates the brain metabolome of SPF

The microbiota can secrete or modulate host secretion of neurotransmitters and neuromodulators implicated in anxiety-like behaviour^1,27,28^. Various microbes have been shown to affect host behaviour directly through small molecules acting on neuroendocrine circuits, and indirectly by influencing the host’s genetic and epigenetic state and immune functions^29–32^. To gain a comprehensive understanding of the factors that drive host metabolism influenced by the microbiota, particularly in the brain by Oligo-MM^12^ commensals, we profiled metabolites in serum and different brain regions of GF mice, Oligo-MM^12^ and SPF mice. Using untargeted metabolomic analysis by liquid chromatography-mass spectrometry, we identified significant differences between mice with microbes (Oligo-MM^12^ and SPF) compared to GF conditions (**Fig. 5A**). In particular, we observed significant differences in metabolites found in the hypothalamus, hippocampus and serum (**Fig. 5A**). The relative abundance of several metabolites was lower in microbiota-colonised mice compared to GF mice, such as proline, succinate, serotonin, nicotinamide adenine dinucleotide phosphate (NADP (+)), phosphoerythronate, hypotaurine, carbomoyl aspartate, pipecolate, 5-hydroxyindole acetate and 2’-deoxyribonucleoside 5’-diphosphate (dUDP) (**Fig. 5B, C**). On the other hand, several metabolites showed higher relative abundance in the hippocampus of microbiota-colonised mice compared to GF mice, such as N-acetyl-isoleucine/leucine (N-Ac-Ile/Leu), spermidine, cysteine sulfate, 3-hydroxybutyrate, kynurenine, 5,6-dihydrouridine, adenylsuccinate, 5-L-glutamyl-L-alanine (GluAla) and L-glutamyl-L-tryptophan (GluTrp) (**Fig. 5B, C**). Interestingly, all of these metabolites (Log2FC>2.0/<-2.0) found in the hippocampus were different in both colonisation models, in SPF and Oligo-MM^12^ mice, compared to GF mice. Only few metabolite were found to be specifically altered only in SPF or only in Oligo-MM^12^ vs. GF mice with the thresholds we used, which is also reflected in the large number of common differentially abundant metabolites between these two groups (**Fig. 5D**). In serum, methionine sulfoxide, quinolinate, lactoylphenylalanine (LacPhe), glutamate/asparagine (GluAsn), hexose-p and lysophosphatidylinositol (LPI-18:00 and LPI-18:01) were among the metabolites with reduced relative abundance in colonised mice compared to GF mice. Instead, several metabolites had higher relative abundance in SPF mice compared to GF mice or in Oligo-MM^12^ mice compared to GF, such as isopentenyl pyrophosphate (isopentenyl-PP), adenosine, adenosine monophosphate (AMP), S-adenosylmethionine (SAM), γ-L-glutamyl-L-cysteine (GluCys), acylcarnitine (AC-(04:1)), guanosine, phosphoserine and cysteine. (**Fig. 5E, F**). Compared to the metabolites found in the hippocampus, the metabolic profile in the serum was more distinct between SPF and Oligo-MM^12^ colonisation, as reflected by the lower number of shared differentially abundant metabolites between these two groups (**Fig. 5D**).

**Fig. 5:**
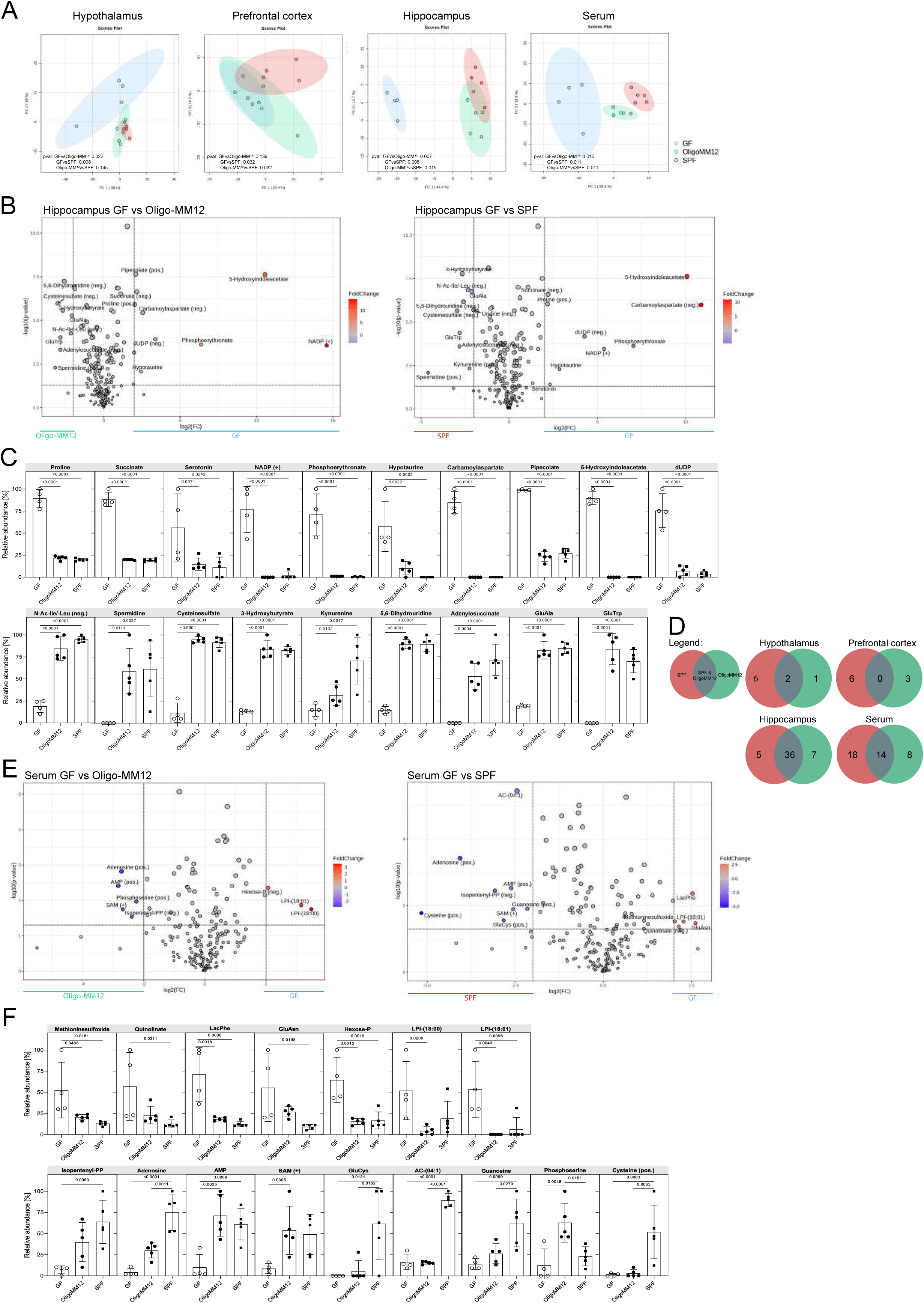
Oligo-MM^12^ microbiota shapes the brain metabolome similarly to SPF microbiota. (**A**) Principal component analysis (PcA) comparing SPF (red), Oligo-MM^12^ (green) and GF (blue) metabolite profile in hypothalamus, prefrontal cortex, hippocampus and the serum. P-values were calculated with pair-wise PERMANOVA tests. Adjusted p-values shown in the plots. (**B**) Volcano plots showing the up-(Log2FC >2.0) /downregulated (< -2.0) metabolites, with the significant ones in blue and red (raw.pval <0.05) in the hippocampus, between GF and Oligo-MM^12^ (left) and GF and SPF (right). (**C**) Relative abundances of the most significantly changed metabolites (Log2FC >2.0/< -2.0, raw.pval <0.05) from (B) in the hippocampus. (**D**) Number of differentially abundant metabolites (raw.pval <0.05, Log2FC >1.0/< -1.0) in colonised vs GF mice, within each brain area and serum. SPF vs. GF only (red), OligoMM12 vs. GF only (green), shared between Oligo-MM^12^ vs. GF & SPF vs. GF (grey). (**E**) Volcano plots showing the up-(Log2FC >2.0) /downregulated (< - 2.0) metabolites, with the significant ones in blue and red (raw.pval <0.05) in the serum, between GF and Oligo-MM^12^ (left) and GF and SPF (right). (**F**) Relative abundances of the most significantly changed metabolites (Log2FC >2.0/< -2.0, raw.pval <0.05) from (E) in the serum. To calculate the relative abundances of the metabolites, peak areas (signal intensities) were normalised to the total metabolite signal intensity of the corresponding sample. The proportion to the highest normalised intensity of the same metabolite across all samples (within one organ) was calculated and plotted as the relative abundance. Sample size: n ≥ 4.

Overall, the differences in host metabolite profiles of moderately diverse microbiota (Oligo-MM^12^) compared to highly diverse microbiota (SPF) are mostly reflected in the serum, whereas in the brain the effects of specific microbial colonisations are more fine-tuned. These data identify metabolites and pathways that may be involved in mediating anxiety-like-behaviour.

## Discussion

The mucosal microbiota significantly influences both healthy and pathological states of host neurology^28,33–41^. From a steady-state neurodevelopmental perspective, animal models have shown that the absence of microbiota is associated with several neurological dysfunctions, such as increased motor activity, exploratory behaviour and home-cage activity and reduced anxiety-like behaviour, as shown by comparing germ-free (GF) with specific pathogen-free (SPF) mice^29^. Furthermore, the depletion of the microbiota in SPF mice using antibiotics facilitates the development of the neurological disorders observed in GF mice. In contrast, colonisation of GF mice with specific commensals and probiotics^1,29,39,42–45^ or prebiotics^46^ can restore the typical behaviour of SPF mice^12,15,47^.

There is also evidence that the gut microbiota can mediate features of neurological diseases^48,49^ and plays a functional role in healthy and unhealthy ageing^50–54^. However, the exact mechanism of action of the microbiota in these contexts is still unknown and we think that it is crucial to study them longitudinally over time. Most studies^1,10–12^ assessing mouse behaviour and the impact of the microbiota on neurological diseases have been conducted only once in the life of the animals, usually as an endpoint and under conventional hygienic conditions where mice are exposed to the environment.

Importantly, changes in the environment, diet and infections shape the microbiota and subsequently affect host physiology, including behaviour^26^. To avoid change and artefacts in the longitudinal study of the effect of the microbiota on behaviour and neurodevelopment, we propose to use gnotobiotic mouse models under controlled hygienic conditions. The sophisticated infrastructure we have established will allow us to bridge the gaps and analyse complex behavioural phenotypes longitudinally in the same animal, subjected to sequential and multiple behavioural tests.

Our study investigates for the first time the effects of Oligo-MM^12^ microbiota on mouse behaviour and shows that Oligo-MM^12^ mice display anxiety-like behaviour similar to SPF mice, in contrast to GF or Cuatro mice. This suggests a major role of the microbiota, and in particular, a very diverse consortia, in anxiety-like behaviour. This is consistent with known differences in anxiety behaviour between different microbiota conditions^12,28,37–41^. Housing conditions also influence behaviour, highlighting the need for precise environmental controls to obtain reproducible results^26^.

The gut-brain axis is influenced by microbial metabolites with neurotrophic effects^12^. We observed similar metabolic profiles in colonised mice, such as Oligo-MM^12^ and SPF mice in different brain regions and serum, and distinct from GF mice (**Fig. 5**). In particular, kynurenine, a metabolite of the tryptophan metabolic pathway, showed higher relative abundance in the hippocampus of Oligo-MM^12^ and SPF mice compared to GF mice, as previously reported^55^. Similarly, 3-hydroxybutyrate, which protects neurons from excitotoxicity and oxidative stress^56,57^, was higher in the hippocampus of Oligo-MM^12^ and SPF mice compared to GF mice. In addition, hippocampal spermidine, a polyamine involved in ameliorating ageing, age-related dementia, and mouse performance in cognitive tests^58^, was positively correlated with *in vivo* colonisation status in our settings. Spermidine is a metabolite from S-adenosylmethionine (SAM) that was detected in the serum of colonised mice in contrast to GF mice (**Fig. 5**). In the circulation, metabolites associated with adenosine production (AMP and adenosine), are enriched in Oligo-MM^12^ and SPF mice compared to GF mice. They have been shown to promote anti-inflammatory and protective functions, including in the gut^59–64^. Adenosine regulates neurotransmitter^65^ release and its production can be enhanced by guanosine, another metabolite present in colonised mice^66–69^. Taken together, our results show for the first time that the Oligo-MM^12^ consortium promotes the production of brain-active and anti-inflammatory molecules, thereby influencing several physiological and neurological functions.

Although our study is limited by the lack of a comparison with SPF mice in isolators and with mice of different ages, sexes, disease stages and health conditions, we think we have provided innovative tools and methods to assess anxiety behaviour and learning and memory functions under controlled hygienic conditions, which are crucial for animal models of neurological disorders such as Alzheimer’s disease. Future research will aim to elucidate the underlying mechanisms that determine the impact of the microbiota on behaviour and neurodevelopment.

## Methods

### Mice

16-22 week old C57BL/6J female germ-free (GF^70^) and gnotobiotic (Cuatro^20^, Oligo-MM^12^ ^4,5^) mice were used. GF mice were routinely monitored for sterility, before and after the behavioural tests, using culture-dependent methods, microbial DNA staining and external laboratory testing. Mice were maintained under axenic conditions in flexible film isolators (NKP Ltd) under a 12 h light/dark cycle. Cuatro and Oligo-MM^12^ were generated by oral gavage of bacteria derived from *in vitro* cultures and have been stably maintained across generations in a series of sterile flexible film isolators. Cuatro microbiota includes four known cultivable commensal bacteria (*Escherichia coli* K12, *Bacteroides thetaiotaomicron* ATCC29148, *Lactobacillus reuteri* I49 and *Clostridium clostridioforme* YL32), while Oligo-MM^12^ consists of 12 known cultivable commensal bacteria^4,5,20^. Stability and absence of contamination in Cuatro and Oligo-MM^12^ consortia, also after behavioural tests (**Fig. S3E, F**), were monitored through routine 16S amplicon sequencing using IonTorrent PGM platform^6^.

20-25 week old C57BL/6JRj female specific-pathogen free (SPF) mice were obtained from Janvier labs (France) and housed at the Center of Phenogenomics (EPFL, Lausanne).

All mouse experiments adhered to Swiss federal regulations approved by the Animal Experimentation Committees of the Cantons of Berne (BE88/17, BE35/21) and Vaud (VD3448).

### Isolator and Mazes at the Clean Mouse Facility of the University of Bern

Behavioural tests were conducted in an M50 isolator (canopy 920 x 1950 x 965 (W x L x H) mm, with a 400 mm internal diameter quadralock) (NKP Ltd). The isolator was prepared, equipped with the material to perform behavioural tests and sterilised with peracetic acid already a week before importing the mice from their original housing isolators. Once transferred to the experimental isolator, the mice were placed in individually ventilated cages (IVC, Tecniplast) and allowed to acclimatise to the new cage and isolator for three days. For each test, the home cage containing the animals to be tested was placed next to the arena to the left of the operator, and the new cage containing the animals after the test was placed to the other side of the arena, to the right of the operator (**Fig. S1A**). In the new cage, soiled bedding and nests from the original home cage were added.

The T-maze apparatus consisted of a two-piece unit (MMTTD-G001, Noldus) made of white acrylic plastic, resistant to peracetic acid (**Fig. S1B**). Three doors (manually operated) were provided to allow individual closing and opening of the 3 arms of the arena. The large open field arena (diameter 630 mm, height 330 mm) is our own innovation developed at the Institute for Surgical Therapies and Biomechanics at the University of Bern, Switzerland, to fit through the isolator’s import door (Quadralock) and the limited working space inside the isolator (**Fig. S1A, S2**). It consisted of a three-piece unit: a circular base, a longitudinal wall panel and a black U-edge strip for the top finish. The base and wall panel were cut from 60 Shore 12mm thick solid white silicone rubber (Silex Ltd UK). The walls were held together with machined stainless-steel screws to form a hairline joint. The edging strip for the top finish was made from 60 Shore 5mm Black Platinum silicone. All parts of the open field arena were designed to be autoclavable at 132°C in a steriliser drum. The small open field arena was custom-made circular in white acrylic plastic (diameter 340 mm, height 560 mm) and sterilised with peracetic acid.

The arenas were placed on a rotating platform (**Figs. S1A**), also an in-house innovation (Institute for Surgical Therapies and Biomechanics at the University of Bern, Switzerland). It consisted of a multi-piece unit consisting of the circular rotary platform itself, a 10 mm melamine-phenolic resin plastic arm and a stainless-steel clamp that held the unit to the Quadralock isolator. The circular rotary platform consisted of two semicircles bent from 10 mm melamine phenolic plastic, which allowed sterilisation at 132°C in a steriliser drum and also entry into the isolator through the 400 mm diameter Quadralock. The plastic arm allowed movement of the circular platform on both the x and y axes to facilitate the operator’s work, as well as movement and focused localisation of the platform and arenas under the camera inside the isolator.

The objects used in the novel object recognition test were rectangular blocks (white or black) 3D printed in Teflon (School of Engineering, EPFL, Lausanne) that can be autoclaved at 132°C in a steriliser drum or sprayed with peracetic acid for sterilisation.

### Behavioural tests at the Clean Mouse Facility of the University of Bern

Animals were housed in groups of five mice per cage and randomised between and within groups during the experiments. The operator was blinded to the hygiene condition throughout the tests. All behavioural tests were performed during the light period between 13:00 and 17:00 h, under dim light at 25 lux. Mouse movements were recorded using a video camera mounted externally on the top of the isolator and analysed using EthoVision XT14 video tracking software (Noldus Information Technology). Our experimental strategy (**Fig. 1A**) was to test each mouse in four different behavioural tests. The arena and objects were cleaned with sterile autoclaved water between each training session, each animal and at the end of each day.

#### Short-Term- and Long-Term Memory tests

Short-Term- and Long-Term Memory tests were conducted in the T-maze adapting a previously published protocol^71,72^. Briefly, both tests consisted of two phases, the habituation phase and the choice phase (**Fig. S1D, E**). The arena was virtually divided into three zones, which were defined by the 3 arms of the T-maze: a starting arm zone, habituation arm zone and a choice arm zone. During the habituation phase (STM/LTM1-4), animals were introduced into the T-maze at the end of the ’start arm’ facing the operator and allowed to freely explore the start arm, central zone and habituation arm of the T-maze, while the choice arm was blocked by an opaque door. Mouse activity was recorded for two minutes. At the end of this time, the mouse was moved to a new cage to rest for one minute for STM or 24 hours for LTM. During the one minute rest period between STM training sessions, the test animal was kept alone in the new cage, whereas during the 24 hour rest period between LTM training sessions, the cage-mates were kept together. After one minute or 24 hours of rest, the mice were reintroduced to the T-maze for a further two minutes of exploration. These steps were repeated for a total of four consecutive exploration periods in the T-maze for the habituation phase with the choice arm always being closed (STM1-4 or LTM1-4). During the subsequent choice phase (STM/LTM choice), the opaque door blocking the third arm (choice arm) was removed and the mice were then reintroduced into the maze for two min and allowed to freely explore the entire surface of the T-maze (**Fig. S1F**).

#### Open Field test (OF)

The open field test was used to test the animals’ anxiety-like behaviour and locomotor activity^73^. Each mouse was introduced into the open field arena in the periphery, on the side of the arena facing the operator. The mouse was free to explore the entire circular arena and its activity was recorded for 20 minutes (**Fig. S3A**). The arena was divided into three virtual circular zones: periphery, which defined a small area along the walls of the arena, a middle zone and the centre zone (**Fig. S3B**). At the end of the exploration period, the animal was moved to a new cage together with all cage mates at the end of their respective sessions.

#### Novel Object Recognition test (NOR)

The Novel Object Recognition test was used to test the animals’ recognition memory for novel environmental cues. The test consisted of three phases: the exploration phase, the learning phase and the recognition phase (**Fig. S3C**). In our experimental design, the 20-minute trial of the open field test also served as the exploration phase of the novel object recognition test. 24 hours after the open field test, the mice were subjected to the learning phase of the NOR. In this phase, two identical objects, in our case white cuboids, were placed in the open field arena. The exact position of the cuboids was determined by measuring the distance from the arena wall in length and width of the object. Each mouse was placed in the periphery of the arena, facing the operator. The mice were free to explore the entire arena and both objects for 10 min. The animal was then moved to a new cage with all cage mates at the end of each session. After three hours of rest, the mice were placed in the arena as described above for the recognition phase in which one of the two objects was replaced by a black cuboid object of same dimensions (novel object). The animal was free to explore the arena with the familiar and novel objects for five minutes. The following virtual zones were defined: familiar object 1 zone and familiar object 1 area zone (a zone surrounding the object) were determined, the same for the other object, the novel object or familiar object 2 (in learning phase) respectively, using the EthoVision XT14 software (**Fig. S3D**).

### Behavioural tests in Lausanne

GF and Oligo-MM^12^ mice bred at the Clean Mouse Facility in Bern were sent to the Center of Phenogenomics in Lausanne (EPFL) seven days before the start of the tests, to acclimatise to the new environment. Before and between the tests, the mice were housed in IVC cages in a conventional animal room with a 12-hour light/12-hour dark cycle. The mice were brought into the experimental room 30 min before the start of the test.

#### Open Field (OF) test

The Open Field and Novel Object Recognition tests were performed as described above, using the same objects used in the tests in Bern. After each session, the mice were returned to their home cage. After each mouse and each day, the arenas were cleaned with 5% ethanol and sterile water. A total of four mice could be tested at a time for the OF test and two mice at a time for the NOR test. The tests were conducted in a sound-attenuated room with dim lighting (18 lux) and isolated from the experimenter. The four arenas (Med Associates) were 400 mm in diameter and (**Fig. S3B, D**) were recorded using an automated video camera system coupled to EthoVision XT14 video tracking software (Noldus Information Technology, The Netherlands). Zones were defined as described above for the tests done in Bern.

### Analysis Behaviour testing data

Each trial was analysed using EthoVision XT14 video tracking software (Noldus Information Technology). For the T-maze exploration, STM and LTM tests, the following parameters were analysed: distance travelled (cm), frequency in each arm (number of entries) and cumulative duration in each arm (s) during the familiarisation and choice phases. In the open field test, distance travelled (cm) and cumulative duration in each zone (s) (zones described in the individual test paragraphs) were quantified. For both these tests, entry into a zone was considered valid when the animal’s midpoint was recorded within the respective zone or arm. For the novel object recognition test, the following parameters were analysed: distance travelled (cm) and the frequency of exploring both objects (number of entries into the object zone with the nose point (facing the object)) in the learning and recognition phases. In this test, a zone entry was considered valid if the animal’s nose point was recorded within the zone.

### Metabolomics

Organs from GF, Oligo-MM^12^ and SPF mice, that were not employed in any behavioural tests, were collected on the same day and under sterile conditions. Blood was taken from the trunk and the serum immediately separated by centrifugation in serum gel tubes (Sarstedt). Hippocampus, hypothalamus and prefrontal cortex were separated from the rest of the brain, snap frozen and stored at -80°C. For metabolite extraction, the frozen samples were carefully thawed on ice. Water-soluble metabolites were extracted from the samples with MeOH/H_2_O (80:20, V:V) containing external standards, diluting the samples in the extraction solution according to their tissue weight and at an organ-specific dilution (1:100 hypothalamus, 1:20 hippocampus, 1:10 prefrontal cortex). Dry sample eluates were redissolved in 100 μL 5 mM NH_4_OAc in acetonitrile/water (50:50, V:V) for tissues, 150 μL for dry serum, and 20 μL of supernatant was loaded into LC vials for LC/MS analysis.

LC-MS analysis was performed on a Thermo Scientific Dionex Ultimate 3000 UHPLC system coupled to a Q Exactive mass spectrometer (QE-MS) equipped with a HESI probe (Thermo Scientific, Bremen, Germany). Mobile phase A was 5 mM NH_4_OAc in acetonitrile/water (50:50, V:V) and mobile phase B was 5 mM NH_4_OAc in acetonitrile/water (95:5, V:V). Measurement of 3 μL injected sample was performed on a BEH Amide column maintained at 45°C with a constant flow rate of 200 μL/min. The initial mobile phase composition was 0:100 (A:B) for 2 min, followed by a linear decrease to 90:10 (A:B) within 23 min, then held constant at 90:10 for 16 min before returning to 0:100 (A:B) in 2 min. 0:100 was maintained for 7 min for column equilibration prior to the next sample injection. The mass spectrometer was set to alternating positive and negative electrospray ionisation (ESI) mode mode and a scan range of 69.0 - 1000 m/z with a resolution of 70,000 was obtained.

Peaks corresponding to the calculated monoisotopic metabolite masses (MIM +/- H+ ± 2 mMU) were integrated using TraceFinder V3.3 software (Thermo Scientific, Bremen, Germany). Peak areas were calculated and normalised to the total metabolite signal intensity of the corresponding sample. Heatmaps, principal component analysis (PcA) and metabolite set enrichment analysis (MSEA) (KEGG) was performed using Metaboanalyst 6.0.

## Figure Legends

**Fig. S1:**
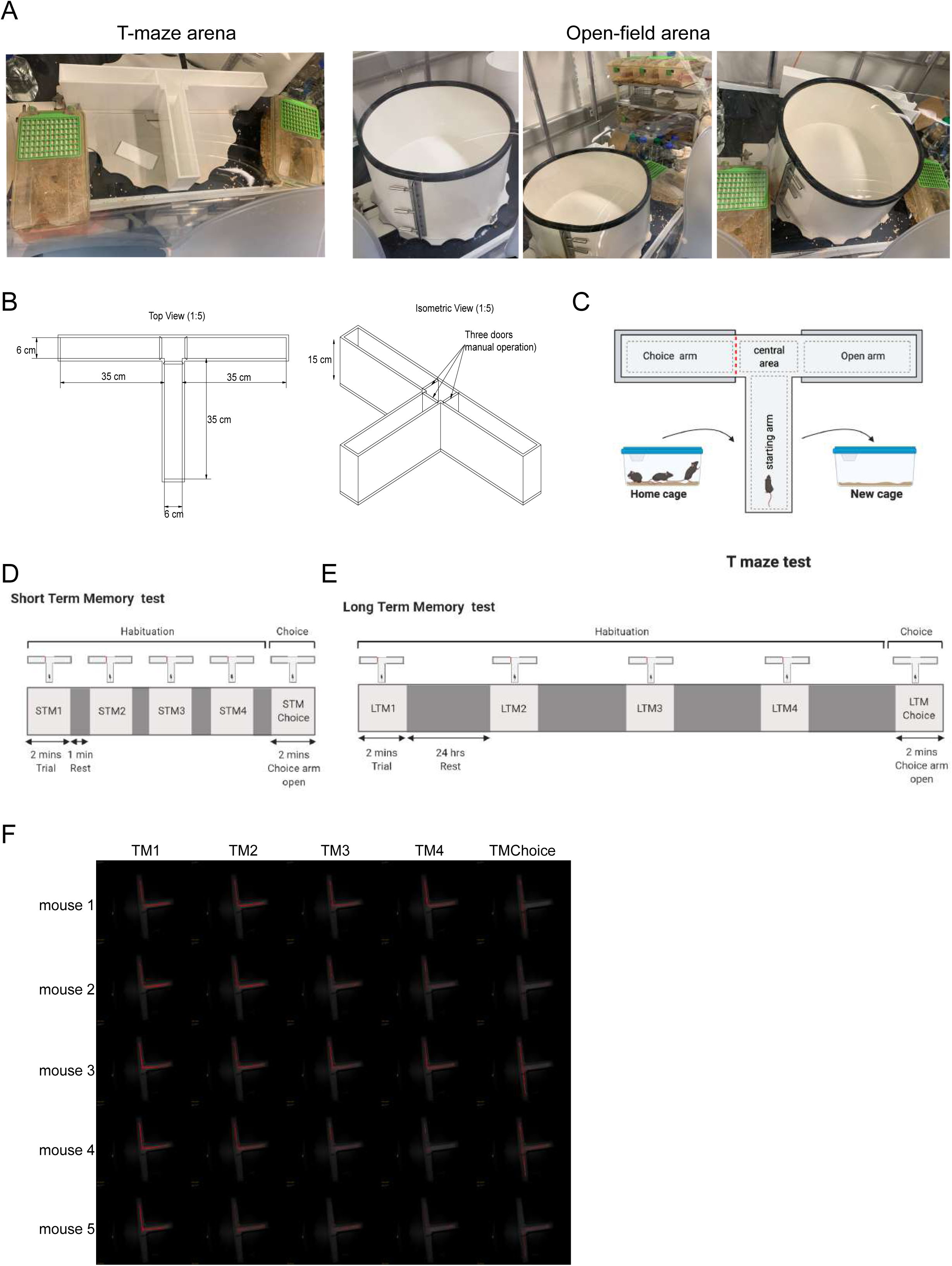
Axenic settings. (**A**) Picture of the settings in flexible film isolators for T-maze arena (left) and pictures of the settings in the flexible film isolator for circular arena (right panels). (**B**) Dimension of the T-maze from top view (left) and isometric view (right). (**C**) Settings of the T-maze test experiments. (**D**) Short-Term Memory protocol. (**E**) Long-Term Memory protocol. (**F**) Representative tracking of the movements of the mice during Short-Term Memory test in the T-maze arena, of different training sessions. Each row is an individual mouse.

**Fig. S2:**
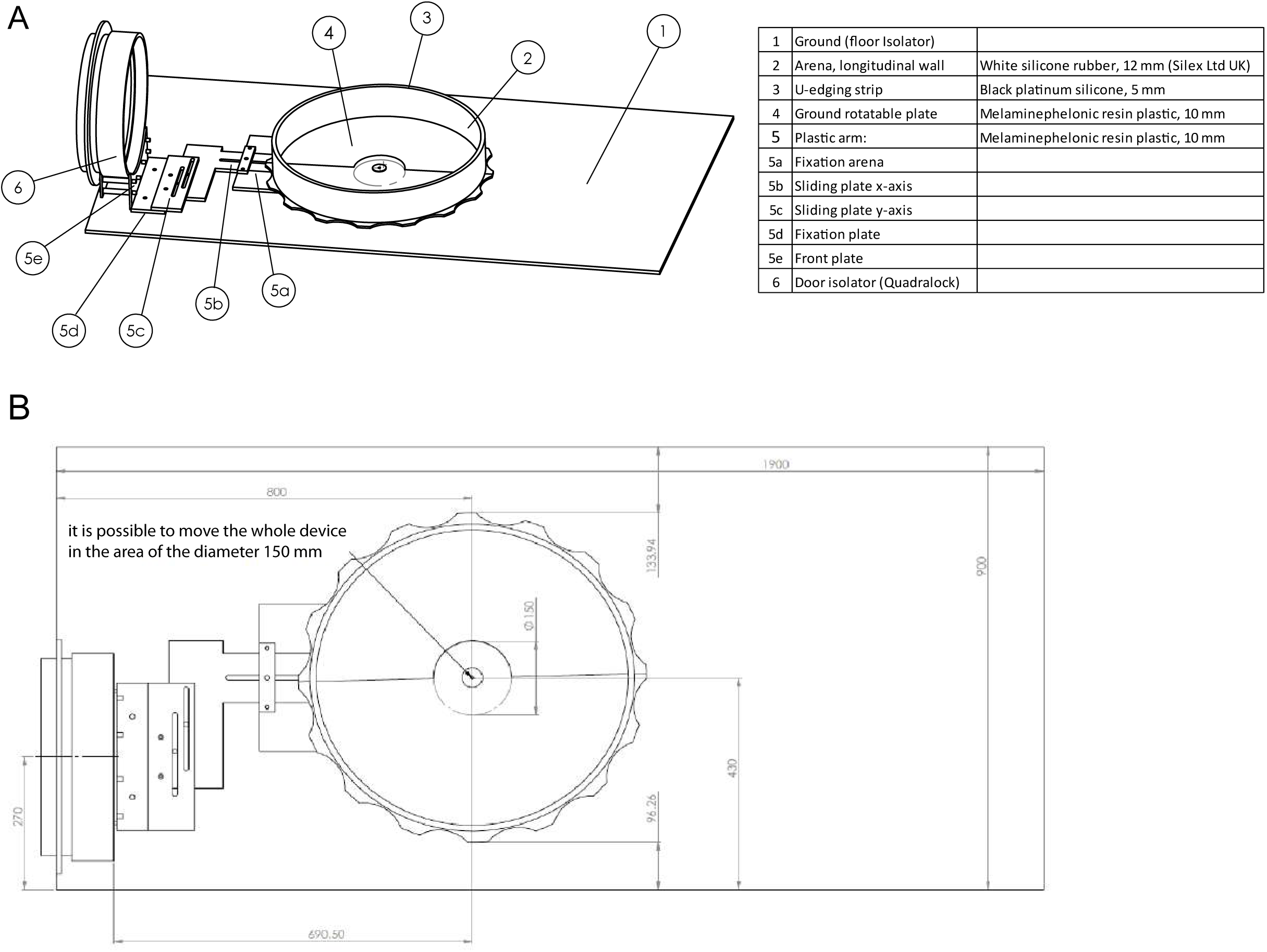
Circular arena set up and experimental set-up in isolators. (**A**) Constituent parts of the circular Open Field arena. (**B**) Top view of the arena with dimensions.

**Fig. S3:**
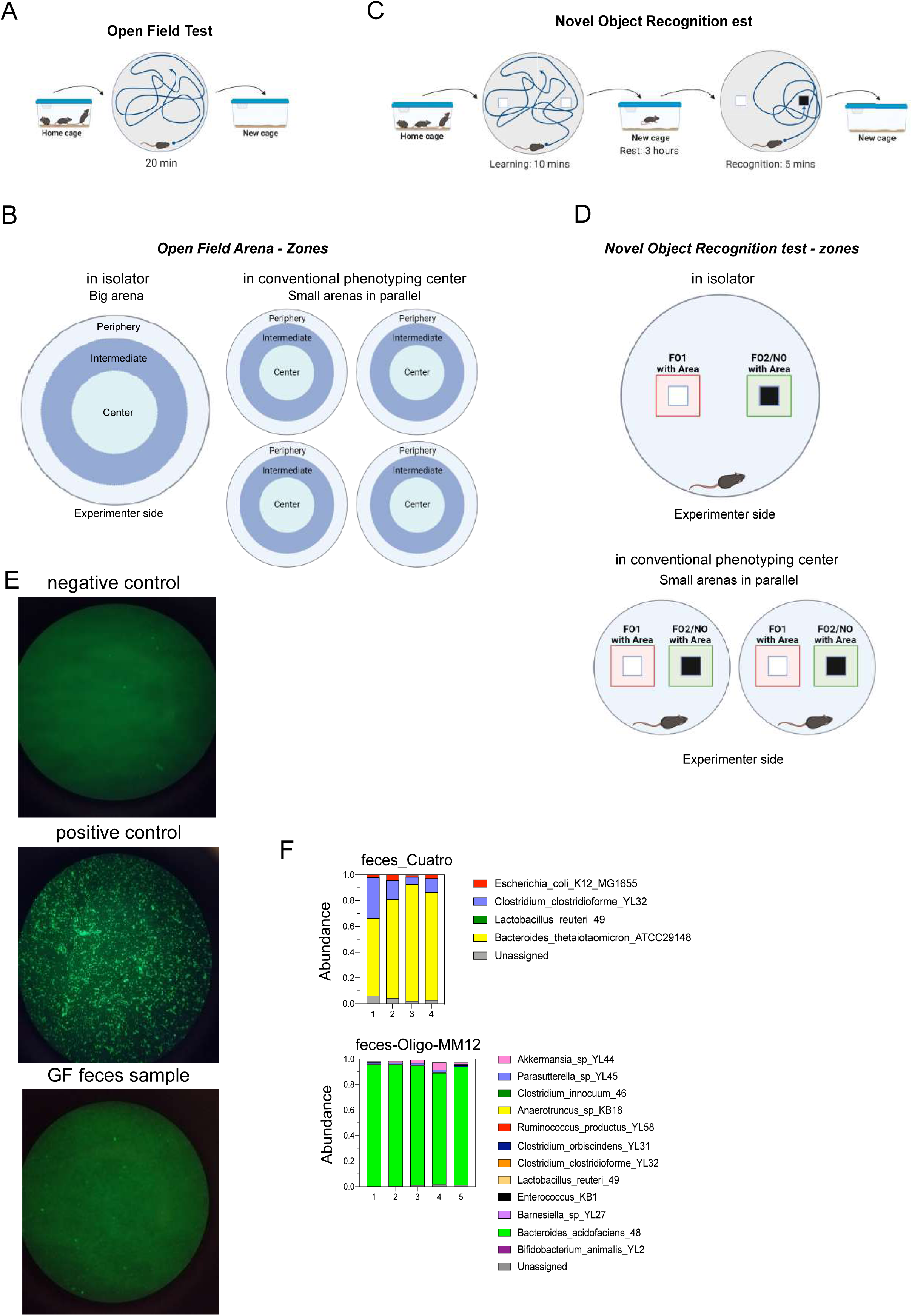
Experimental settings for Open Field, Novel Object Recognition and Morris Water Maze tests. (**A**) Experimental plan for Open field test. (**B**) Arena settings with the different defined zones of the Open Field test in Bern (left) and in Lausanne (right). (**C**) Experimental plan for Novel Object Recognition test. (**D**) Settings of the Novel Object Recognition test in Bern (top) and Lausanne (low). (**E**) Sytox staining on faeces of GF mice post-behavioural tests. Negative control is sterile Phosphate Buffered Saline (PBS), positive control is *E.coli* culture. (**F**) Microbial composition in feces at the species level in Cuatro and Oligo-MM^12^-colonised mice post-behavioural tests. Each bar represents a single mouse. Unassigned are sequences that do not map on bacterial databases.

## Acknowledgments

We would like to thank Dr Camin Dean (The German Center for Neurodegenerative Diseases, Berlin) and Melissa Long (Animal Outcome Core Facility / Animal Behavior Phenotyping Facility, ExzellenzCluster NeuroCure, Charité Universitätsmedizin) for critical input into the manuscript. We are grateful for support by the Clean Mouse Facility which is supported by the Genaxen Foundation, Inselspital and the University of Bern and to Alessio Mylonas (EPFL).

## Author contributions

AR, MI, MGA and FR performed all experiments and analysed data. JK, JBK, ALP, TM helped performing experiments. AR, MI, JK, MGA, and FR set up all experimental conditions in gnotobiotic hygiene settings. WS performed and analysed the metabolomics analysis. RD helped in designing the behavioural tests. MGA and FR designed the study and experiments and wrote the manuscript. All authors revised the manuscript and approved its final version.

